# Automation and Evaluation of the SOWH Test with SOWHAT

**DOI:** 10.1101/005264

**Authors:** Samuel H. Church, Joseph F. Ryan, Casey W. Dunn

## Abstract

The Swofford-Olsen-Waddell-Hillis (SOWH) test evaluates statistical support for incongruent phylogenetic topologies. It is commonly applied to determine if the maximum likelihood tree in a phylogenetic analysis is significantly different than an alternative hypothesis. The SOWH test compares the observed difference in likelihood between two topologies to a null distribution of differences in likelihood generated by parametric resampling. The test is a well-established phylogenetic method for topology testing, but is is sensitive to model misspecification, it is computationally burdensome to perform, and its implementation requires the investigator to make multiple decisions that each have the potential to affect the outcome of the test. We analyzed the effects of multiple factors using seven datasets to which the SOWH test was previously applied. These factors include bootstrap sample size, likelihood software, the introduction of gaps to simulated data, the use of distinct models of evolution for data simulation and likelihood inference, and a suggested test correction wherein an unresolved “zero-constrained” tree is used to simulate sequence data. In order to facilitate these analyses and future applications of the SOWH test, we wrote SOWHAT, a program that automates the SOWH test. We find that inadequate bootstrap sampling can change the outcome of the SOWH test. The results also show that using a zero-constrained tree for data simulation can result in a wider null distribution and higher p-values, but does not change the outcome of the SOWH test for most datasets. These results will help others implement and evaluate the SOWH test and allow us to provide recommendation for future applications of the SOWH test. SOWHAT is available for download from https://github.com/josephryan/SOWHAT.

---

A phylogenetic topology test evaluates whether the difference in optimality criterion score between incongruent hypotheses is significant. In some cases, the test is used to determine whether a dataset provides significantly more support for one of several previously proposed phylogenetic hypotheses. In other cases, the test is used to compare a novel or unexpected phylogenetic result to a previously proposed hypothesis. In both scenarios, the observed difference in the optimality criterion score between two trees is compared to an estimated null distribution of differences in scores. Phylogenetic topology tests differ largely in how this null distribution is created. The Kishino-Hasegawa test (KH) (Kishino and Hasegawa 1989), for example, creates a null distribution by analyzing datasets created by sampling with replacements from the original dataset (Goldman et al. 2000). However, this approach is subject to selection bias and is only appropriate for tests of hypotheses selected *a priori*, such as the comparison of two alternative hypotheses from the literature. In cases where hypotheses are not determined *a priori*, as when comparing a topology to an unanticipated maximum likelihood tree produced from the same data, tests such as the Approximately Unbiased test (AU) (Shimodaira 2001), the Shimodaira-Hasegawa test (SH) (Shimodaira and Hasegawa 1999), and the Swofford-Olsen-Waddell-Hillis test (SOWH) are more appropriate (Goldman et al. 2000). The SOWH test, as a parametric bootstrap method, can provide greater statistical power than non-parametric methods such as the SH test, though this comes at the cost of an increased reliance on the model of evolution (Goldman et al. 2000).

A typical SOWH test where an *a priori* hypothesis is compared to the maximum likelihood tree (Fig. 1) begins with performing two maximum likelihood searches on a dataset. One is unconstrained and results in the most likely tree. The other is constrained by an alternative hypothesis, which will result in a less likely topology. The difference in likelihood scores between these two topologies (*δ*) is the test statistic. New datasets are then simulated using the topology and parameter estimates (base frequencies, rates, alpha value, etc) from the constrained likelihood search. On each of these datasets, two maximum likelihood searches - one unconstrained and one constrained by the alternative hypothesis - are performed and *δ* values are calculated. The test statistic *δ* is then compared to this distribution of simulated *δ* values. A significant test statistic is one which falls above some percent, such as 95%, of simulated *δ* values.

**Figure 1:**
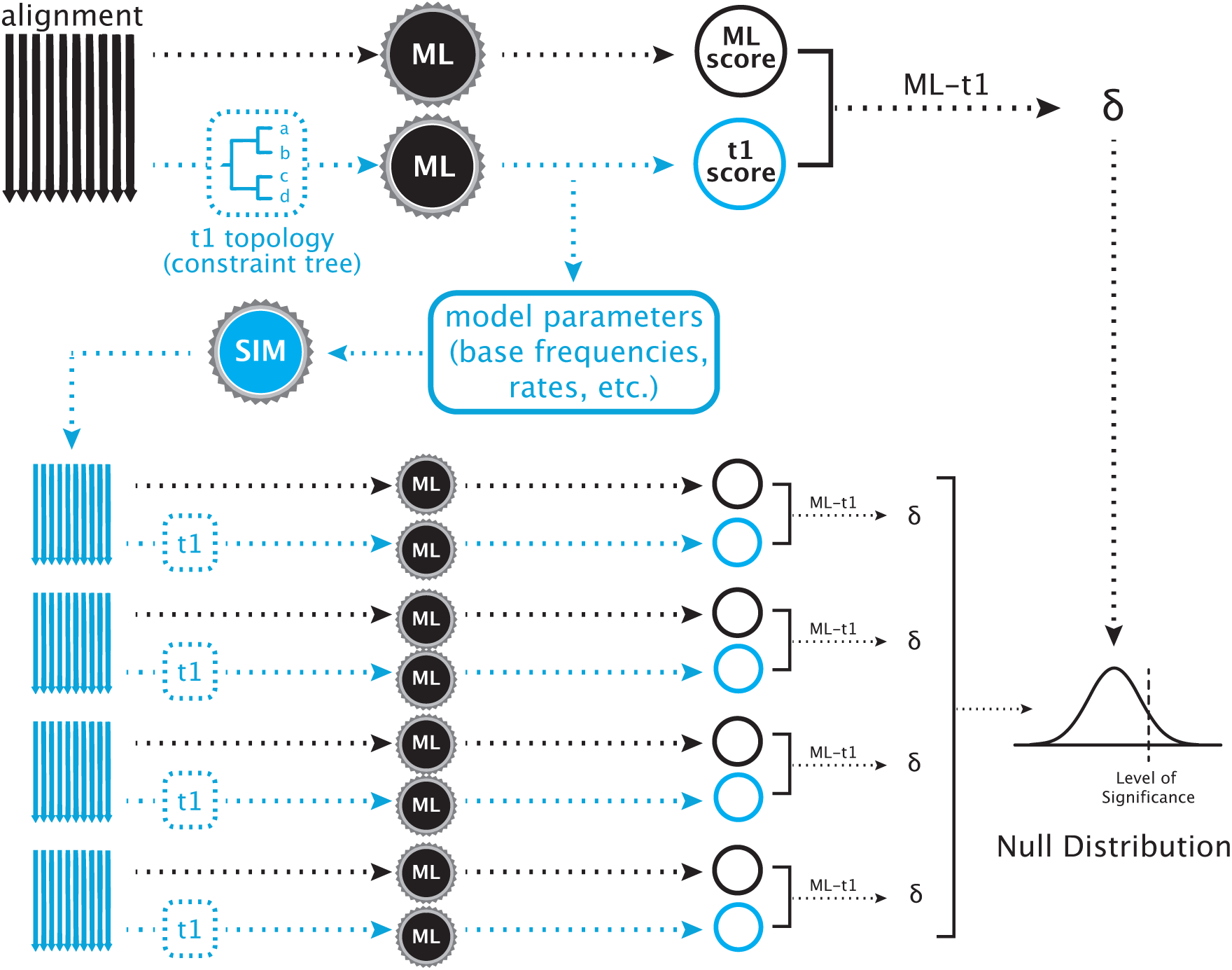
**A Typical SOWH Test** The test begins with two maximum likelihood searches on a single alignment. One search, represented by the black arrow, is performed with no constraining topology. Another test, represented by the blue arrow, is constrained to follow an *a priori* topology that represents a phylogenetic hypothesis incongruent with the maximum likelihood topology. The black gears represent maximum likelihood software used to score the trees (i.e. GARLI, RAxML). These two searches result in two maximum likelihood scores, the difference (*δ*) between which is the test statistic. From the constrained search, the optimized parameters and topology are retrieved and used to simulate new alignments with software (blue gear) such as Seq-Gen. For each simulated alignment (blue), two maximum likelihood searches are performed, one unconstrained (black arrow) and one constrained (blue arrow), scores are obtained, and a *δ* value is calculated. The test statistic is compared to this distribution of *δ* values. A significantly large *δ* value is one which falls above some proportion of those generated by data simulation (i.e. 95%)

The SOWH test can be burdensome to implement with existing tools. Several helpful step-by-step instructions for manual implementation are available (Crawford 2009; Anderson et al. 2014), but these approaches require extensive hands-on time which makes it difficult to systematically examine the behavior of the test under different conditions. Performing the SOWH test requires an investigator to make multiple decisions which may not be informed without an evaluation of the behavior of the test. These decisions include how many bootstrap samples to generate, which likelihood software to use, and how to treat gaps during data simulation. These factors regularly vary across SOWH tests published in the literature and prior to the analysis described here there has not been an evaluation of their effects on multiple datasets.

Furthermore, the test has been shown to have a high type I error rate as compared to other topology tests (Buckley 2002; Susko 2014). In order to correct this, two adjustments to the implementation of the SOWH test have been recommended. Prior to the present study, these hypothesized corrections have not been evaluated across multiple datasets. Buckley (2002) suggested that when the model of evolution for simulating data is the same model used in a maximum likelihood search on the data, it is more probable that the generating topology will be found. Because the generating topology in a SOWH test conforms to the alternative hypothesis, both the constrained and unconstrained searches on the simulated data will find a tree very similar to the generating tree. This will result in smaller *δ* values, a more narrow null distribution, and a potentially higher rate of type I error. To correct for this, previous investigators recommended that the model of evolution for likelihood inference should be distinct from the model for simulating data (Buckley 2002; Pauly et al. 2004) (Fig. 2).

**Figure 2:**
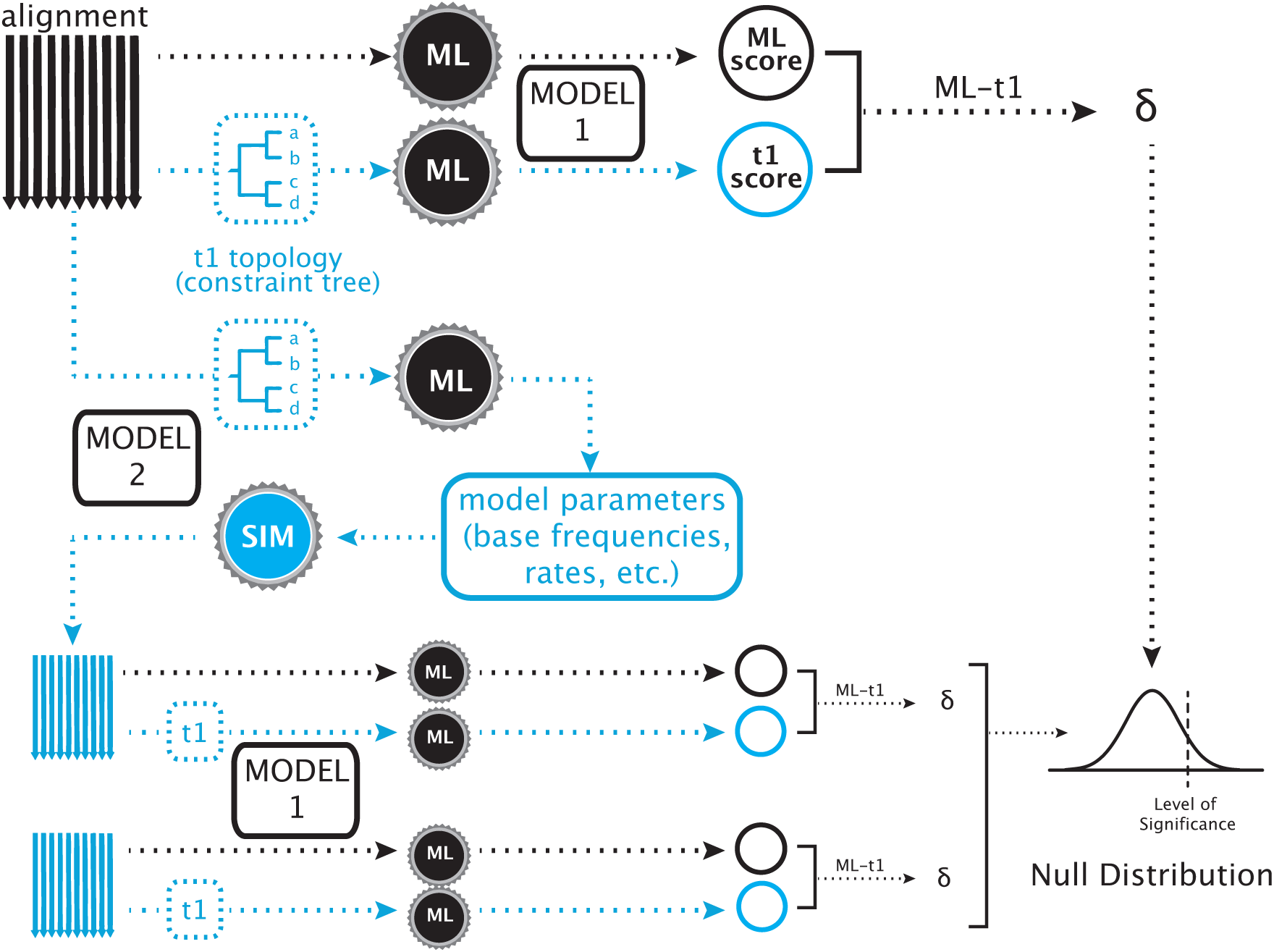
**A SOWH Test Using Two Models of Evolution** In this test, two maximum likelihood searches are performed as described above using a model of evolution (Model 1). Instead of retrieving parameter values from the constrained search, as would be done in a typical SOWH test, an additional constrained maximum likelihood is performed using a different model of evolution (Model 2). Parameters are retrieved from this test and used to simulate new data. These simulated datasets are then scored using the same model of evolution used to score the original datasets (Model 1). This adjustment to the typical SOWH test was suggested following the assumption that a SOWH test performed with the same model of evolution for likelihood scoring and data simulation would result in a smaller *δ* values on simulated data, a smaller null distribution, and a more liberal test. For tests in our study which use the CAT-GTR model in PhyloBayes, both the additional constrained search and data simulation are performed using PhyloBayes - all other likelihood searches are performed using the specified likelihood software (i.e. GARLI or RAxML).

More recently Susko (2014) observed that likelihood differences between unconstrained and constrained searches on simulated data are smaller than would be expected. Susko (2014) suggests that when data are simulated on a fully resolved tree, it is more likely that the generating topology used for simulation will be found. This will result in a more narrow distribution and a potentially higher type I error rate, as described above. As a correction, Susko (2014) suggests using a tree which is not fully resolved to simulate data, specifically the newly proposed “zero-constraineded” tree. This tree is generated by manipulating the most likely tree estimated in an unconstrained search so that all edges incongruent with the alternative hypothesis are reduced to zero or near zero, creating a polytomy.

Using seven published datasets previously analyzed with the SOWH test, we examined the effects on test outcome of multiple factors (sample size, likelihood software, and treatment of gaps) and proposed adjustments (model specification and generating topology). To facilitate implementation, we developed SOWHAT (as in, “The maximum likelihood tree differs from my alternative phylogenetic hypothesis, so what?”), a program that automates the SOWH test (Fig. 1) and includes features that allow for additional complexities such as partitioned datasets. Using our results we provide recommendations for future implementation of the SOWH test. These recommendations, along with the program SOWHAT, allow for a more informed and less burdensome application of the SOWH test.

## METHODS AND MATERIALS

### Implementation of the SOWH test with SOWHAT

We used SOWHAT, our automation of the SOWH test, for all analyses. SOWHAT is available at https://github.com/josephryan/SOWHAT.

This tool evaluates the significance of the difference between the unconstrained maximum likelihood tree and a maximum likelihood tree inferred under a topology constraint provided by the user (Fig. 1). At a minimum, the user specifies the model of evolution under which the maximum likelihood searches will be performed, and provides as input an alignment file (in PHYLIP format) as well as the constraint topology to be tested (in Newick format). Two maximum likelihood trees - an unconstrained tree, and a tree constrained according to the provided constraint topology - are then inferred with GARLI (Zwickl 2006) or RAxML (Stamatakis 2006), as specified by the user.

New datasets are simulated by Seq-Gen (Rambaut and Grass 1997), or by PhyloBayes (Lartillot et al. 2009) if the CAT-GTR model is selected for simulation. The simulated alignments are generated using the topology, branch lengths, and model parameters (i.e., state frequencies, rates, and the alpha parameter for the gamma rate heterogeneity approximation) from the constrained analysis as inferred by the likelihood software. If the correction of Susko (2014) is applied, the data are instead simulated on the zero-constrained tree. If the original dataset is partitioned, parameters are estimated separately for each partition and the alignments are generated following the partitioning scheme. Each of the simulated alignments is then evaluated using the likelihood software, again using both an unconstrained search and a constrained search. The difference in likelihood of these two searches (*δ*) is calculated for each simulated dataset, and the set of these differences make up the null distribution.

The original likelihood difference (*δ*) is then compared against the null distribution using a Monte-Carlo method, given by the equation *p* = *d/n* where *d* is the number of *δ* values greater than or equal to the test statistic and *n* is the sample size.

SOWHAT can be used to evaluate a hypothesized topology given datasets of nucleotide, amino acid, or binary characters. All models for likelihood inference available in RAxML or GARLI are available in SOWHAT.

### Calculation of Confidence Intervals

SOWHAT calculates confidence intervals around a p-value following the addition of each new sample to the null distribution. The confidence intervals are calculated using a normal approximation interval, given by the equation 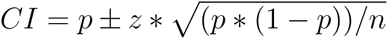, where p is the proportion of total samples greater than the test statistic and z is the z-value from the normal distribution (Ellison and Gotelli 2004). SOWHAT calculates the confidence intervals assuming a significance level of 95%, therefore *z* = 1.96. This approximation is only valid at a sufficiently large sample size. The confidence values can be used to determine adequacy of sample size by evaluating whether both the lower and upper interval fall on the same side of the significance level, 0.05.

### Examined Datasets

We used Google Scholar (http://scholar.google.com) to perform a literature search for papers that cited Goldman et al. (2000) and included the terms “SOWH” or “parametric bootstrap”. We narrowed these roughly 400 results down to 40 which had deposited data on TreeBase (http://treebase.org/). From these we selected seven datasets for examination that represent a range of p-values and dataset sizes. They are included with SOWHAT so that users can easily reproduce our analyses. The results presented here were prepared with the version of SOWHAT (v0.20) available at https://github.com/josephryan/sowhat/commit/3137601014e24274ebc115acabe086290b5f46e7. We analyzed the data with GARLI version 2.01.1067, RAxML version 8.1.15, PhyloBayes version 3.3f, seq-gen version 1.3.2x, and R version 3.0.0.

*Buckley (2002).*— From the analysis of the SOWH test by Buckley (2002), we selected a dataset of mitochondrial ribosomal RNA genes (12S) originally assembled by Sullivan et al. (1995). Likelihood analysis of this alignment produces a topology of sigmodontine rodents, which differs from the topology recovered by analyses of morphological, chromosomal, allozyme, and other DNA datasets (Buckley 2002). Buckley (2002) performed a series of SOWH tests using PAUP* and multiple models, all of which rejected the alternative hypothesis using a significance level of 0.05. Here we repeat the analyses using the models GTR+I+Γ and GTR+Γ. The dataset contains eight taxa and 791 characters.

*Dixon et al. (2007).*— This dataset contains 14 individuals of five species of European *Androsace*, a perennial herbaceous plant (Dixon et al. 2007). The alternative hypothesis to the maximum likelihood topology includes a monophyletic haplotype group of chloroplast regions for two species, *A. laggeri* and *A. pyrenica*. Dixon et al. (2007) performed a SOWH test using PAUP* which rejected this hypothesis using the HKY model of evolution, but not when using HKY+Γ. The dataset contains 2180 nucleotide characters.

*Dunn et al. (2005).*— This dataset is composed of mitochondrial ribosmal RNA genes (16S and 18S) from 52 siphonophores (Hydrozoa: Cnidaria), as well as four outgroup taxa (Dunn et al. 2005). The authors tested several hypotheses - the one we examine here unites *Bargmannia* with Agalmatidae *sensu stricto*. Dunn et al. (2005) performed a SOWH test which rejected this hypothesis, using parsimony (PAUP*) to score the original and simulated trees and maximum likelihood to estimate parameters and simulate data. The dataset contains 2748 nucleotide characters.

*Edwards et al. (2005).*— This dataset is composed of five distinct gene regions from 38 taxa in the group Cactaceae (Edwards et al. 2005). The alternative hypothesis to the maximum likelihood topology creates a monophyletic group, *Pereskia*. Edwards et al. (2005) performed a SOWH test which rejected this hypothesis, using PAUP* (ML) and the model GTR+I+Γ. The dataset contains 6150 nucleotide characters.

*Liu et al. (2012).*— This dataset includes 12 DNA loci, including 3 nuclear loci, 5 chloroplast loci, and 4 mitochondrial loci from 26 species of bryophytes (Liu et al. 2012). The alternative hypothesis to the maximum likelihood topology constrains the monophyly of *Physcomitrella*. Liu et al. (2012) performed a SOWH test which rejected this hypothesis using RAxML and the model GTR+I+Γ, and incorporated a partitioning scheme which we did not utilize in the analyses presented here. The dataset contains 44 taxa and 6657 nucleotide characters.

*Sullivan et al. (2000).*— This dataset is composed of mitochondrial sequences from 26 individuals of the rodent species *Reithrodontomys sumichrasti*. The alternative hypothesis to the maximum likelihood topology constrains phylogeographic groups similar to other rodents (Sullivan et al. 2000). Sullivan et al. (2000) performed SOWH tests using both maximum likelihood (under a GTR+I+Γ model) and parsimony optimality criterion scores in PAUP*. The hypothesis was rejected using maximum likelihood scores but not using parsimony scores. The dataset contains 1130 nucleotide characters.

*Wang et al. (2008).*— This dataset is composed of mitochondrial sequences from 41 individuals of the species of pygmy rain frog *Pristimantis ridens* and a single outgroup taxon Wang et al. (2008). The alternative hypothesis incongruent with the maximum likelihood constrains a phylogeographic of seven individuals. Wang et al. (2008) performed a SOWH test which rejected this hypothesis using PAUP* and fixed parameter values for maximum likelihood searches of simulated datasets. The dataset contains and 1672 nucleotide characters.

### Sensitivity to Bootstrap Sample Size

We selected three of the seven datasets for an analysis of sensitivity to sample size. These datasets were selected because the results reported in the original study did not strongly reject the alternative hypothesis. For each of these datasets we performed 100 SOWH tests using a sample size of 100 each, and then subsequently performed 100 SOWH tests using a sample size of 500 each. We compared the range of p-values returned to the average confidence intervals calculated by SOWHAT at the two sample sizes. Each of these tests was performed using RAxML and the model GTR+Γ.

### Sensitivity to Choice of Likelihood Software

For all seven datasets we performed a test using GARLI and a test using RAxML, with the exception of Dixon et al. (2007) because the model HKY is not an option in RAxML. We compared the results of five of these tests to the originally reported results which were performed using PAUP*(ML). Our analyses were performed using the same model, sample size, and constraint topology as reported in the literature. For our analyses, gaps were propagated into simulated data - treatment of gaps was mentioned only in the study by Edwards et al. (2005) where gaps were not present in simulated data. It is not clear how gaps were treated in the remaining datasets. We did not directly compare the results for Dunn et al. (2005), because the original study used parsimony to score datasets, or Liu et al. (2012), because the original study used a partitioning scheme not used for these analyses. For all datasets except Dixon et al. (2007), we also compared the results of the tests performed using GARLI to those performed using RAxML. Following consultation with the authors of RAxML, we reran the RAxML analyses with the BFGS routine suppressed (using the --nobfgs flag in RAxML) and compared the results.

### Sensitivity to Treatment of Gaps

By default, SOWHAT propagates the same number and position of gaps present in the original dataset into each simulated dataset. This is accomplished by simulating a full dataset and then subsequently removing the characters in matrix positions corresponding to gaps in the original dataset. We analyzed the effect of not including gaps by suppressing this feature (i.e. simulated data matrices were complete, where originals were not) and comparing the resulting p-values to this calculated with gaps in simulated data. Four of the seven datasets analyzed in this study had gaps in the original data (Dunn et al. (2005), Edwards et al. (2005), Liu et al. (2012), and Wang et al. (2008)). We used RAxML as the likelihood engine in all of these searches, and the same sample size as reported in the original study.

### Sensitivity to Model Specification

We analyzed the effects of using a different model for likelihood inference from the model used for parameter estimation and data simulation (Fig. 2). For all seven datasets we performed a SOWH test in which the likelihood score was calculated using a model with a reduced number of parameters free to vary (JC69) in comparison to the model used for parameter estimation and data simulation (GTR+I+Γ). We compared the resulting p-values to those calculated using the originally reported model of evolution for both inference and data simulation. For these tests we used GARLI as the likelihood engine and the same sample size as reported in the original study. We then ran a SOWH test in which the originally reported model of evolution was used exclusively for likelihood inference and a model with an increased number of free parameters (the CAT-GTR model found in PhyloBayes) was used for parameter estimation and data simulation. The resulting p-values were compared to those calculated using the originally reported model for both likelihood and inference. For these tests we used RAxML as the likelihood engine and the same sample size as reported in the original study.

### Sensitivity to Generating Topology

We analyzed the effects of simulating data under the zero-constrained tree in place of a fully resolved tree estimated under the alternative hypothesis. In this method, following calculation of the most likely tree in an unconstrained search, SOWHAT uses the R package hutan (https://bitbucket.org/caseywdunn/hutan) to identify incongruent nodes and reduce them to an epsilon value near zero (here 1 * 10^−6^). All other branch lengths are preserved, resulting in a partially resolved tree referred to as the zero-constrained tree (Susko 2014). For all seven datasets we compared the results of a SOWH test using a fully resolved generating topology to a test using the zero-constrained tree. We used RAxML as the likelihood engine in all of these searches, with the exception of Dixon et al. (2007) where GARLI was used, and the same sample size as reported in the original study.

## RESULTS AND DISCUSSION

### Bootstrap Sample Size

When an investigator applies the SOWH test they must decide how many simulated bootstrap samples to generate. Without sampling an adequate number of replicates, stochasticity can lead to poor estimates of the null distribution and different outcomes between repeated SOWH tests. Increasing the number of replicates, however, comes with an increased computational cost. Few previously reported SOWH tests provide justification for sample size. A sufficient sample size depends on the data in question. In order to explore the effects of sample size, we ran multiple identical SOWH tests and compared the resulting p-values.

Our results indicate that a sample size of 100 is not sufficient for all datasets (Table 1). For the Dixon et al. (2007) dataset, repeated SOWH tests with 100 samples returned p-values ranging from 0.030 to 0.160, which results in a different interpretation of outcome at a significance level of 0.05. When the sample size was increased to 500, the test consistently returned a p-value greater than 0.05.

**Table 1:**
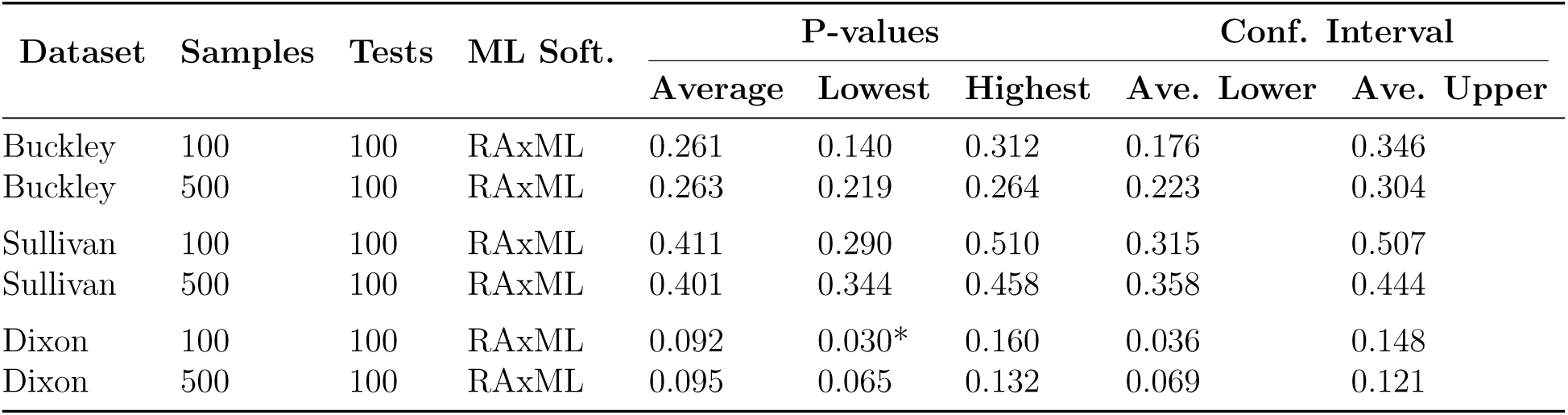
**Bootstrap Sample Size** 100 SOWH tests were performed for three datasets with a sample size of 100, and 100 tests were performed with a sample size of 500. P-values for the Dixon et al. (2007) dataset at a sample size of 100 vary from 0.030 to 0.169, indicating repeated SOWH tests at this sample size could result in different outcomes using a significance level of 0.05. The average confidence interval ranges from 0.036 to 0.148, indicating that confidence intervals capture a large range of observed variation and that they could be used to evaluate the adequacy of sample size. At a sample size of 500 all p-values and the average confidence intervals fell above the level of significance. * indicates p-value less than 0.05.

The confidence intervals calculated by SOWHAT provide an explicit tool for assessing the adequacy of sampling for a given test. With each new value added to the null distribution, SOWHAT recalculates the p-value and confidence intervals, allowing the adequacy of the sample size to be evaluated simultaneously with the results of the test. In the case of the SOWH tests on Dixon et al. (2007) with 100 samples, the average confidence interval spans from 0.036 to 0.148, which includes the threshold of 0.05. This signals to the investigator that there is not adequate sampling to provide a robust interpretation of the outcome. As sample size increases, the confidence interval decreases, and at a sample size of 500 the average lower confidence interval for this dataset falls above the level of significance. This indicates that sample size is sufficient.

### Choice of Likelihood Software

Application of the SOWH test also requires deciding which software to use for likelihood estimation. Our results indicate that, all else being equal, the choice of likelihood software between GARLI and RAxML does not affect the outcome of the SOWH test as long as a minor adjustment is made to how RAxML is run. Specifically, BFGS optimization of RAxML model parameters is inconsistent on simulated data, but suppressing this optimization resolves the problem.

We compared the results of SOWH tests performed using the same data, model, and sample size, but different likelihood software (GARLI and RAxML) for six datasets (Table 2). For the Buckley (2002) we performed this comparison using two different models for each likelihood software tool. Under BFGS optimization (the RAxML default), the choice of likelihood software had an effect on the outcome for two datasets. The SOWH test of Sullivan et al. (2000) performed with GARLI rejected the hypothesis, where the SOWH test performed using RAxML failed to reject the hypothesis (P *>* 0.05). As well, the SOWH test of Wang et al. (2008) performed with GARLI rejected the hypothesis, where the SOWH test using RAxML failed to reject the hypothesis (P *>* 0.01).

**Table 2:** **Choice of Likelihood Software** SOWH tests were performed using GARLI and RAxML for each dataset, with the exception of Dixon et al. 2007 as HKY is not an option in RAxML. Outcomes of the tests were different between likelihood programs for analyses of Sullivan et al. (2000) and Wang et al. (2008). Each SOWH test was performed using the model, sample size, and constraint topology specified in the original performance of the test. Buckley (2002) and Dixon et al. (2007) were analyzed using two different models of evolution, as reported originally. The resulting p-values were compared to those reported in the literature for five datasets. The other two datasets were not directly compared due to known differences in implementation; the SOWH test performed by Dunn et al. (2005) used parsimony to score tree; the test by Liu et al. (2012) was performed with a partition scheme not used here. The outcome of the tests differed from the literature for two additional datasets, Buckley (2002) using GTR+Γ and Dixon et al. (2007) using HKY. * indicates p-values less than 0.05; ** indicates less than 0.01.

| Dataset | Source | Samples | Model | ML Software | P-value | Conf. Interval |  |
| --- | --- | --- | --- | --- | --- | --- | --- |
|  |  |  |  |  |  | Lower | Upper |
| Buckley | New | 1000 | GTR+I+$\Gamma$ | GARLI | 0.018* | 0.010 | 0.026 |
| Buckley | New | 1000 | GTR+I+$\Gamma$ | RAxML | 0.025* | 0.015 | 0.035 |
| Buckley | Reported | 1000 | GTR+I+$\Gamma$ | PAUP* (ML) | 0.018* | - | - |
| Buckley | New | 1000 | GTR+$\Gamma$ | GARLI | 0.118 | 0.098 | 0.138 |
| Buckley | New | 1000 | GTR+$\Gamma$ | RAxML | 0.241 | 0.215 | 0.268 |
| Buckley | Reported | 1000 | GTR+$\Gamma$ | PAUP* (ML) | 0.015* | - | - |
| Dixon | New | 500 | HKY | GARLI | 0.012* | 0.002 | 0.021 |
| Dixon | Reported | 500 | HKY | PAUP* (ML) | 0.258 | - | - |
| Dixon | New | 500 | HKY+$\Gamma$ | GARLI | 0.002** | -0.002 | 0.005 |
| Dixon | Reported | 500 | HKY+$\Gamma$ | PAUP* (ML) | <0.001** | - | - |
| Dunn | New | 100 | GTR+I+$\Gamma$ | GARLI | <0.001** | <0.001 | <0.001 |
| Dunn | New | 100 | GTR+I+$\Gamma$ | RAxML | <0.001** | <0.001 | <0.001 |
| Edwards | New | 100 | GTR+I+$\Gamma$ | GARLI | <0.001** | <0.001 | <0.001 |
| Edwards | New | 100 | GTR+I+$\Gamma$ | RAxML | <0.001** | <0.001 | <0.001 |
| Edwards | Reported | 100 | GTR+I+$\Gamma$ | PAUP* (ML) | <0.01** | - | - |
| Liu | New | 100 | GTR+I+$\Gamma$ | GARLI | <0.001** | <0.001 | <0.001 |
| Liu | New | 100 | GTR+I+$\Gamma$ | RAxML | <0.001** | <0.001 | <0.001 |
| Sullivan | New | 100 | GTR+I+$\Gamma$ | GARLI | <0.001** | <0.001 | <0.001 |
| Sullivan | New | 100 | GTR+I+$\Gamma$ | RAxML | 0.290 | 0.201 | 0.379 |
| Sullivan | Reported | 100 | GTR+I+$\Gamma$ | PAUP* (ML) | <0.01** | - | - |
| Wang | New | 500 | GTR+$\Gamma$ | GARLI | <0.001** | <0.001 | <0.001 |
| Wang | New | 500 | GTR+$\Gamma$ | RAxML | 0.026* | 0.012 | 0.040 |
| Wang | Reported | 500 | GTR+$\Gamma$ | PAUP* (ML) | <0.005** | - | - |

In order to understand the underlying cause of these different outcomes, we compared the null distributions and test statistics generated by each of the SOWH tests performed using GARLI and RAxML (Table 3). We did not observe differences in the calculated test statistics on the real data between RAxML and GARLI that would change the outcome of the test, suggesting that this component of the analysis is not responsible for the discrepancies. Initial results instead indicated that the differences in significance between programs are due to irregularities in RAxML performance on the simulated data used to generate the null distribution. For every SOWH test performed, the range of the null distribution generated using RAxML was larger than the range generated using GARLI. This is because RAxML occasionally returned large *δ* values in both tails of the distribution relative to those returned by GARLI. The presence of these large values in the left tail of the null distribution signifies that for certain simulated datasets RAxML returns a much lower likelihood score using an unconstrained search than using a constrained search. This indicated that RAxML was not identifying the most likely tree in the unconstrained searches on simulated data, as the constrained topology is within the treespace of the unconstrained search. Following consultation with the authors of RAxML, the problem was identified as a failure of the BFGS optimization of model parameters. Rerunning the tests using RAxML with the optimization suppressed resulted in universally smaller ranges of null distribution (Table 3) and the outcomes of the tests run under RAxML were the same as those under GARLI.

**Table 3:** **Effects of Likelihood Software on Null Distributions** A SOWH test was performed using GARLI and RAxML for each dataset. RAxML initially resulted in a larger total range of the null distribution for all datasets. Large values in the left tail of the distribution, as seen in the Sullivan et al. (2000) and Wang et al. (2008) analyses, indicate failure of RAxML to find the most likely topology. Following consultation with the authors of RAxML, a SOWH test was performed using RAxML with the BFGS routine suppressed for each dataset, which resulted in a smaller range in all cases and no extreme values in the tails. The outcomes of the tests between GARLI and RAxML with the BFGS routine suppressed are equivalent. Test statistics varied between SOWH tests, though this variance alone would not have had an effect on the final outcome.

| Dataset | Model | ML Software | P-value | Test Stat. | Null Distribution |  |  |  |  |  |
| --- | --- | --- | --- | --- | --- | --- | --- | --- | --- | --- |
|  |  |  |  |  | 0% | 25% | 50% | 75% | 100% | Range |
| Buckley | GTR+I+$\Gamma$ | GARLI | 0.018* | 4.703 | -0.147 | 0.102 | 0.831 | 1.775 | 8.022 | 8.168 |
| Buckley | GTR+I+$\Gamma$ | RAxML | 0.025* | 4.875 | -55.879 | <0.001 | 0.010 | 1.118 | 73.743 | 129.622 |
| Buckley | GTR+I+$\Gamma$ | RAxML (No BFGS) | 0.010* | 4.873 | -1.076 | 0.000 | 0.534 | 1.489 | 9.530 | 10.605 |
| Buckley | GTR+$\Gamma$ | GARLI | 0.118 | 4.888 | -0.860 | 0.165 | 1.348 | 3.254 | 17.192 | 18.052 |
| Buckley | GTR+$\Gamma$ | RAxML | 0.241 | 4.961 | -21.136 | 0.214 | 2.385 | 4.787 | 28.437 | 49.573 |
| Buckley | GTR+$\Gamma$ | RAxML (No BFGS) | 0.252 | 4.871 | -1.581 | 0.499 | 2.566 | 4.873 | 23.381 | 24.962 |
| Dunn | GTR+I+$\Gamma$ | GARLI | <0.001** | 21.796 | -0.023 | <0.001 | 0.279 | 1.101 | 4.499 | 4.522 |
| Dunn | GTR+I+$\Gamma$ | RAxML | <0.001** | 17.810 | -4.319 | <0.001 | 0.767 | 2.808 | 8.592 | 12.911 |
| Dunn | GTR+I+$\Gamma$ | RAxML (No BFGS) | <0.001** | 16.427 | -6.292 | 0.015 | 0.707 | 2.062 | 5.276 | 11.569 |
| Edwards | GTR+I+$\Gamma$ | GARLI | <0.001** | 9.684 | -0.001 | <0.001 | <0.001 | <0.001 | 0.008 | 0.009 |
| Edwards | GTR+I+$\Gamma$ | RAxML | <0.001** | 9.732 | -0.039 | -0.003 | -0.002 | <0.001 | 0.019 | 0.058 |
| Edwards | GTR+I+$\Gamma$ | RAxML (No BFGS) | <0.001** | 9.731 | -0.032601 | -0.004 | -0.002 | <0.001 | 0.009 | 0.041 |
| Liu | GTR+I+$\Gamma$ | GARLI | <0.001** | 335.653 | -0.001 | <0.001 | <0.001 | 0.298 | 4.009 | 4.010 |
| Liu | GTR+I+$\Gamma$ | RAxML | <0.001** | 335.620 | -0.041 | -0.001 | 0.002 | 0.094 | 4.018 | 4.059 |
| Liu | GTR+I+$\Gamma$ | RAxML (No BFGS) | <0.001** | 335.628 | -1.805 | 0.000 | 0.002 | 0.304 | 4.831 | 6.637 |
| Sullivan | GTR+I+$\Gamma$ | GARLI | <0.001** | 8.828 | <0.001 | 0.320 | 1.354 | 2.720 | 7.434 | 7.434 |
| Sullivan | GTR+I+$\Gamma$ | RAxML | 0.290 | 8.413 | -560.827 | 0.612 | 2.139 | 10.575 | 581.588 | 1,142.414 |
| Sullivan | GTR+I+$\Gamma$ | RAxML (No BFGS) | <0.001** | 8.466 | -1.308 | 0.155 | 0.730 | 1.957 | 7.049 | 8.357 |
| Wang | GTR+$\Gamma$ | GARLI | <0.001** | 12.542 | -0.002 | 0.000 | 0.200 | 1.169 | 8.442 | 8.444 |
| Wang | GTR+$\Gamma$ | RAxML | 0.026* | 13.018 | -652.832 | 0.012 | 0.269 | 1.515 | 780.203 | 1,433.035 |
| Wang | GTR+$\Gamma$ | RAxML (No BFGS) | <0.001** | 13.026 | -2.802 | 0.005 | 0.190 | 1.262 | 10.337 | 13.139 |

We also compared the results of the SOWH test reported in the literature to the results obtained here for five of these seven datasets. Each of the tests compared was performed using the same hypothesis, model, and sample size as originally reported, but differed in likelihood software - all of the reported tests were previously performed using PAUP*. In addition to the previously stated difference in results of the Sullivan et al. (2000) and Wang et al. (2008) tests, two other tests returned outcomes different than those previously reported. For Buckley (2002) using the model GTR+Γ the SOWH tests performed using GARLI and RAxML both failed to reject a hypothesis reported to be rejected in the original study (P *<* 0.05). For Dixon et al. (2007) the SOWH test performed using GARLI and the model HKY rejected a hypothesis (P *<* 0.05) that was not rejected in the original study. As the reported SOWH tests were not performed using SOWHAT, we cannot determine whether these results are due to differences in differences in performance of likelihood software or other aspects of test implementation. We did not directly compare the results of the Dunn et al. (2005) or Liu et al. (2012) due to known differences in implementation beyond likelihood software - despite these difference, for both datasets the outcome of the SOWH tests performed here were consistent with those originally reported.

### Treatment of Gaps

Multiple sequence alignments include gaps due to both missing data and the accommodation of insertions and deletions. These gaps are often not included in the simulated bootstrap samples. The purpose of simulating datasets under parametric conditions is to recreate the situation under which the real-world data was generated - ideally this would recreate the presence of gaps as well.

SOWHAT simulates the data without gaps and subsequently replaces determined sites with gaps based on the number and positions of gaps in the original dataset. While this does not explicitly model the processes that led to the original gaps, this method ensures that taxa which were originally poorly sampled and which may be in question will be poorly sampled in all simulated datasets as well.

We tested the effects of gaps by suppressing this feature (i.e. all simulated datasets were completely full). Our results indicate that, for the four datasets tested here, gaps had no major effect on the outcome of the SOWH test (Table 4). Excluding these gaps from simulated data did not change the outcome of the test for any of these datasets.

**Table 4:** **Treatment of Gaps** SOWHAT by default propagates the exact number and position of gaps present in the original dataset into all simulated datasets. Suppressing this feature, thereby excluding gaps from subsequent analyses, did not change the outcome of any SOWH tests examined here. * indicates p-values less than 0.05; ** indicates less than 0.01.

| Dataset | % gaps in dataset | Simulate Gaps | P-value | Conf. Interval |  |
| --- | --- | --- | --- | --- | --- |
|  |  |  |  | Lower | Upper |
| Dunn | 14.759 | yes | <0.001** | <0.001 | <0.001 |
| Dunn | 14.759 | no | <0.001** | <0.001 | <0.001 |
| Edwards | 14.268 | yes | <0.001** | <0.001 | <0.001 |
| Edwards | 14.268 | no | <0.001** | <0.001 | <0.001 |
| Liu | 9.149 | yes | <0.001** | <0.001 | <0.001 |
| Liu | 9.149 | no | <0.001** | <0.001 | <0.001 |
| Wang | 13.092 | yes | 0.030* | 0.015 | 0.045 |
| Wang | 13.092 | no | 0.032* | 0.017 | 0.047 |

### Model Specification

The choice of model used for likelihood inference and for simulating datasets has previously been shown to have an impact on the outcome of the SOWH test (Goldman et al. 2000; Buckley 2002). In order to minimize type I errors associated with model misspecification, Buckley (2002) suggested that the model used for likelihood inference should be different from the model used to estimate parameters and simulate data (Fig. 2). They suggest that when the likelihood search is being performed with the exact same model used to generate the data it will be prone to recover the generating topology, therefore *δ* values will be smaller and the null distribution will be more narrow.

We evaluated the effect of this adjustment by implementing a simpler model for likelihood inference than the model used for data simulation, using two different methods. First, for seven datasets we compared the results of SOWH tests performed using the JC69 model for likelihood inference and GTR+I+Γ for data simulation (Table 5) to tests using the same model for likelihood inference and data simulation. Our results show that, contrary to what was previously suspected, using distinct models resulted in lower *δ* values and a potentially higher type I error rate. For every test performed using JC69 for likelihood simulation the p-value returned was *<*0.001 and the hypothesis was rejected, including tests which did not reject the hypothesis when the same model was used for likelihood inference and data simulation.

**Table 5:** **Model Specification: JC69 Analysis** Model 1 represents the model used for likelihood inference (i.e. searching and scoring both the original and simulated datasets). Model 2 was used to calculate parameter values and simulate datasets. Separating the models for scoring and simulation has been suggested as a correction for a liberal bias present in the SOWH test. Using a model for scoring with fewer parameters free to vary, such as JC69, here resulted in a more liberal test. All hypotheses were rejected when JC69 was used as Model 1. * indicates p-values less than 0.05; ** indicates less than 0.01.

| Dataset | Models |  | P-value | Conf. Interval |  |
| --- | --- | --- | --- | --- | --- |
|  | ML Score | Parameters |  | Lower | Upper |
| Buckley | GTR+G | GTR+G | 0.241 | 0.214 | 0.268 |
| Buckley | JC69 | GTR+G+I | <0.001** | <0.001 | <0.001 |
| Dixon | HKY+G | HKY+G | 0.012* | 0.002 | 0.022 |
| Dixon | JC69 | GTR+G+I | <0.001** | <0.001 | <0.001 |
| Dunn | GTR+G+I | GTR+G+I | <0.001** | <0.001 | <0.001 |
| Dunn | JC69 | GTR+G+I | <0.001** | <0.001 | <0.001 |
| Edwards | GTR+G+I | GTR+G+I | <0.001** | <0.001 | <0.001 |
| Edwards | JC69 | GTR+G+I | <0.001** | <0.001 | <0.001 |
| Liu | GTR+G+I | GTR+G+I | <0.001** | <0.001 | <0.001 |
| Liu | JC69 | GTR+G+I | <0.001** | <0.001 | <0.001 |
| Sullivan | GTR+G+I | GTR+G+I | 0.290 | 0.201 | 0.379 |
| Sullivan | JC69 | GTR+G+I | <0.001** | <0.001 | <0.001 |
| Wang | GTR+G+I | GTR+G+I | 0.026* | 0.012 | 0.040 |
| Wang | JC69 | GTR+G+I | <0.001** | <0.001 | <0.001 |

Second, for six datasets we also compared the results of SOWH tests when parameters were optimized and data simulated under the PhyloBayes CAT-GTR model and a simpler model used for inference (Table 6). Our results show that this method can change the outcome of the test in both directions, depending on the data in question. For Buckley (2002) and Sullivan et al. (2000), the SOWH test performed using the CAT-GTR model rejected a hypothesis which was not rejected using the same model for inference and simulation (P *>* 0.05). Conversely, for Dunn et al. (2005) the SOWH test performed using the CAT-GTR model failed to reject a hypothesis which was rejected using the same model for inference and simulation (P *<* 0.05). These results do not support the hypothesis that separating the models will universally result in higher *δ* values and a more conservative test. We find no consistent evidence to support the suggestion that the model for inference should be distinct from the model for simulation.

**Table 6:** **Model Specification: CAT Analysis** Model 1 and Model 2 are the same as described in Table 5. Using a model for simulation with a greater number of parameters free to vary, such as the CAT model of PhyloBayes, did not result in universally larger *δ* values and therefore a more conservative test, though this was true for one dataset, Dunn et al. (2005). The outcome of two other tests also differed, for Buckley (2002) and Sullivan et al. (2000), but the result was a more conservative test. * indicates p-values less than 0.05; ** indicates less than 0.01.

| Dataset | Models |  | P-value | Conf. Interval |  |
| --- | --- | --- | --- | --- | --- |
|  | ML Score | Parameters |  | Lower | Upper |
| Buckley | GTR+$\Gamma$ | GTR+$\Gamma$ | 0.241 | 0.214 | 0.268 |
| Buckley | GTR+$\Gamma$ | CAT | 0.013* | 0.006 | 0.020 |
| Dunn | GTR+I+$\Gamma$ | GTR+I+$\Gamma$ | <0.001** | <0.001 | <0.001 |
| Dunn | GTR+I+$\Gamma$ | CAT | 0.080 | 0.026 | 0.133 |
| Edwards | GTR+I+$\Gamma$ | GTR+I+$\Gamma$ | <0.001** | <0.001 | <0.001 |
| Edwards | GTR+I+$\Gamma$ | CAT | <0.001** | <0.001 | <0.001 |
| Sullivan | GTR+I+$\Gamma$ | GTR+I+$\Gamma$ | 0.290 | 0.201 | 0.379 |
| Sullivan | GTR+I+$\Gamma$ | CAT | 0.030* | 0.003 | 0.063 |
| Liu | GTR+I+$\Gamma$ | GTR+I+$\Gamma$ | <0.001** | <0.001 | <0.001 |
| Liu | GTR+I+$\Gamma$ | CAT | <0.001** | <0.001 | <0.001 |
| Wang | GTR+G | GTR+G | 0.026* | 0.012 | 0.040 |
| Wang | GTR+G | CAT | 0.020* | 0.008 | 0.032 |

**Table 7:**
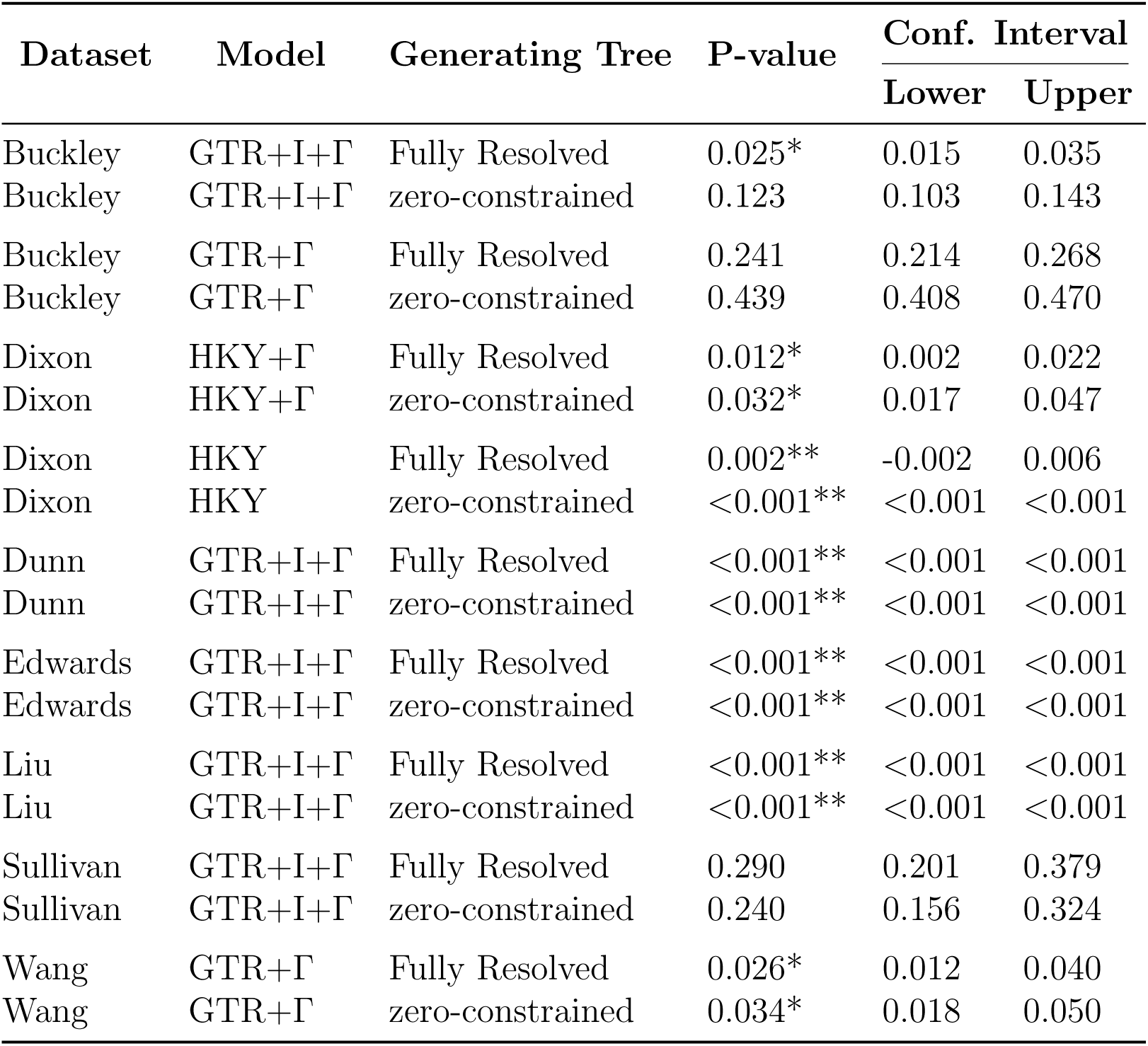
**Generating Topology** We compared SOWH tests performed using a fully resolved generating topology to tests performed using the zero-constrained tree, as suggested by Susko (2014). The zero-constrained tree is created by manipulating the most likely unconstrained tree so that edges incongruent with the alternative hypothesis are reduced to nearly zero. Using this method changed the outcome of only one test, Buckley (2002), using the model GTR+I+Γ. * indicates p-values less than 0.05; ** indicates less than 0.01.

It has also been suggested that, in order to reduce the rate of type I error, parsimony might be used in place of likelihood inference to score the datasets (Buckley 2002). Parsimony has often been used to reduce the computational burden of running multiple likelihood searches (Dunn et al. 2005). However using parsimony in the context of the SOWH test is problematic. Often the purpose of the test is to determine whether the most likely hypothesis significantly more likely than some *a priori* hypothesis. It is very possible that the *a priori* hypothesis may not be significantly less parsimonious and still be significantly less likely, as seems to be the case in the two SOWH tests reported in Sullivan et al. (2000). Given the improved speed in likelihood software, parsimony should no longer be used in place of likelihood in the context of the SOWH test.

### Generating Topology

The SOWH test, as presented in Goldman et al (2000), generates new datasets using the parameter values and tree topology estimated under the alternative constraint hypothesis. Susko (2014) suggest that using a fully resolved tree as the generating topology will result in a test that is too conservative and that returns lower p-values than it should. This suggestion is based on the hypothesis that a likelihood search performed on simulated data generated under a fully resolved tree will more easily find the generating topology. As the generating topology used in the SOWH test conforms to the constrained alternative hypothesis, both the constrained and unconstrained searches will therefore find the same generating topology, and the *δ* values will be close to zero.

Susko (2014) suggests instead using a third tree, referred to as the zero-constrained tree, as the generating topology. This tree is created by manipulating the most likely tree estimated in an unconstrained search so that all edges incongruent with the alternative hypothesis are reduced to zero or near zero. The resulting topology preserves all branch lengths and nodes that are not incongruent with the constraint tree resulting in a partially resolved tree. Reducing incongruent edges results in a polytomy for the nodes in question. Susko (2014) hypothesis that this method will result in larger *δ* values and therefore a less conservative test.

We evaluated this adjustment by comparing the results of SOWH tests performed using a fully resolved tree to the results of tests performed using the zero-constrained tree for all seven datasets included in this study. Our results show that using the zero-constrained tree results in very similar p-values for nearly all of the tests in question. The outcome of the test was different for one datasets, Buckley (2002) when using the model GTR+I+Γ, where the test using the zero-constrained tree did not reject the alternative hypothesis which was rejected using a fully resolved generating tree (P *>* 0.05).

These results suggest that using the zero-constrained tree can result in higher p-values, but that the impact is minimal and for many datasets the outcome of the test is unchanged.

## CONCLUSION

The SOWH test is of great value to the phylogenetic community. However, it can be burdensome to perform manually and requires making multiple decisions that have the potential to affect the outcome of the test. SOWHAT eliminates the burden of manual implementation of the SOWH test. Using SOWHAT to evaluate the impact of these decisions, we provide the following recommendations for implementation of the SOWH test. Where possible, these recommendations have been adopted as the default behavior of SOWHAT, which makes comparing results across SOWH tests easier.

The SOWH bootstrap sample size should be explicitly justified. Our results indicate that a sample size of 100 is not adequate for all datasets. SOWHAT calculates confidence intervals around the p-values, and our results indicate that these confidence values capture a reasonable amount of the observed variation in repeated tests. Following a minimum number of samples - we recommend 100 - an investigator should evaluate the adequacy of sample size by determining whether the confidence intervals fall entirely on one side of the line of significance.

We found that the choice of likelihood software did not depend on whether GARLI or RAxML were used for likelihood analyses, as long as BFGS optimization was suppressed in RAxML. Given that RAxML performs the likelihood searches much faster than GARLI, RAxML with the BFGS routine silened is the default likelihood engine in SOWHAT. GARLI can be used if additional models are required.

Our results indicate that excluding gaps from simulated data is unlikely to change the overall outcome of a SOWH test, however this effect may depend on the data in question. Since there is little computational cost and strong biological justification, we recommend simulating data with gaps.

Model misspecification is a problem universal to parametric bootstrap tests, which rely on the model to both simulate and evaluate the data. It has been suggested that using a model for likelihood inference that is distinct from the model used for data simulation may result in lower type I error rate Buckley (2002). Our results are not consistent with this hypothesis - we found that using a simpler model for inference compared to the model used for simulation does not result in universally higher *δ* values. We therefore suggest that the same model is used for inference and simulation.

It has also been suggested that simulating new data under a tree which is not fully resolved, such as the zero-constrained tree will result in lower type I error Susko (2014). We find that simulating data on the zero-constrained tree can impact results, but that for many datasets using the zero-constrained tree will not change the outcome of the test. Given the well-established concern of a high type I error rate along with the low computation cost in implementation, we recommend simulating data under the zero-constrained tree.

## ACKNOWLEDGMENTS

SHC was funded by the LINK Award through Brown University and the Marine Biological Laboratory at Woods Hole, as well as the Sars International Centre for Marine Molecular Biology. JFR was funded by The Whitney Laboratory for Marine Bioscience and Sars International Centre for Marine Molecular Biology. We thank Nicholas Sinnott-Armstrong, Felipe Zapata, Mark Howison, Peisi Yan, Stefan Siebert, Andreas Hejnol for comments, ideas, and support. Analyses were conducted with computational resources and services at the Center for Computation and Visualization at Brown University, supported in part by the NSF EPSCoR EPS-1004057 and the State of Rhode Island. Those acknowledged do not necessarily endorse the findings in this paper.

## Reference

Anderson, J., N. Goldman, and A. Rodrigo. 2014. Guidelines for performing the sowh test http://www.ebi.ac.uk/goldman/tests/sowhinstr.html.

Buckley, T. R. 2002. Model misspecification and probabilistic tests of topology: evidence from empirical data sets. Systematic Biology 51:509–523.

Crawford, A. J. 2009. horribly detailed instructions to running a parametric bootstrap test http://dna.ac/genetics.html.

Dixon, C. J., P. Schoenswetter, and G. M. Schneeweiss. 2007. Traces of ancient range shifts in a mountain plant group (androsace halleri complex, primulaceae). Molecular ecology 16:3890–3901.

Dunn, C. W., P. R. Pugh, and S. H. Haddock. 2005. Molecular phylogenetics of the siphonophora (cnidaria), with implications for the evolution of functional specialization. Systematic Biology 54:916–935.

Edwards, E. J., R. Nyffeler, and M. J. Donoghue. 2005. Basal cactus phylogeny: implications of pereskia (cactaceae) paraphyly for the transition to the cactus life form. American Journal of Botany 92:1177–1188.

Ellison, G. N. and N. Gotelli. 2004. A primer of ecological statistics. Sinauer, Sunderland, Massachusetts, USA.

Goldman, N., J. P. Anderson, and A. G. Rodrigo. 2000. Likelihood-based tests of topologies in phylogenetics. Systematic Biology 49:652–670.

Kishino, H. and M. Hasegawa. 1989. Evaluation of the maximum likelihood estimate of the evolutionary tree topologies from dna sequence data, and the branching order in hominoidea. Journal of molecular evolution 29:170–179.

Lartillot, N., T. Lepage, and S. Blanquart. 2009. Phylobayes 3: a bayesian software package for phylogenetic reconstruction and molecular dating. Bioinformatics 25:2286–2288.

Liu, Y., J. M. Budke, and B. Goffinet. 2012. Phylogenetic inference rejects sporophyte based classification of the funariaceae (bryophyta): rapid radiation suggests rampant homoplasy in sporophyte evolution. Molecular phylogenetics and evolution 62:130–145.

Pauly, G. B., D. M. Hillis, and D. C. Cannatella. 2004. The history of a nearctic colonization: molecular phylogenetics and biogeography of the nearctic toads (bufo). Evolution 58:2517–2535.

Rambaut, A. and N. C. Grass. 1997. Seq-gen: an application for the monte carlo simulation of dna sequence evolution along phylogenetic trees. Computer applications in the biosciences: CABIOS 13:235–238.

Shimodaira, H. 2001. Multiple comparisons of log-likelihoods and combining nonnested models with applications to phylogenetic tree selection. Communications in Statistics-Theory and Methods 30:1751–1772.

Shimodaira, H. and M. Hasegawa. 1999. Multiple comparisons of log-likelihoods with applications to phylogenetic inference. Molecular biology and evolution 16:1114–1116.

Stamatakis, A. 2006. Raxml-vi-hpc: maximum likelihood-based phylogenetic analyses with thousands of taxa and mixed models. Bioinformatics 22:2688–2690.

Sullivan, J., E. Arellano, and D. S. Rogers. 2000. Comparative phylogeography of mesoamerican highland rodents: concerted versus independent response to past climatic fluctuations. The American Naturalist 155:755–768.

Sullivan, J., K. E. Holsinger, and C. Simon. 1995. Among-site rate variation and phylogenetic analysis of 12s rrna in sigmodontine rodents. Molecular Biology and Evolution 12:988–1001.

Susko, E. 2014. Tests for two trees using likelihood methods. Molecular biology and evolution Page msu039.

Wang, I. J., A. J. Crawford, and E. Bermingham. 2008. Phylogeography of the pygmy rain frog (pristimantis ridens) across the lowland wet forests of isthmian central america. Molecular phylogenetics and evolution 47:992–1004.

Zwickl, D. 2006. Garli: genetic algorithm for rapid likelihood inference. See http://www.bio.utexas.edu/faculty/antisense/garli/Garli.html.

